# Oligogenic inheritance of congenital heart disease involving a NKX2-5 modifier

**DOI:** 10.1101/266726

**Authors:** Casey A. Gifford, Sanjeev S. Ranade, Ryan Samarakoon, Hazel T. Salunga, T. Yvanka de Soysa, Yu Huang, Ping Zhou, Aryé Elfenbein, Stacia K. Wyman, Yen Kim Bui, Kimberly R. Cordes Metzler, Philip Ursell, Kathryn N. Ivey, Deepak Srivastava

## Abstract

Complex genetic inheritance is thought to underlie many human diseases, yet experimental proof of this model has been elusive. Here, we show that a human congenital heart defect, left ventricular non-compaction (LVNC), can be caused by a combination of rare, inherited heterozygous missense single nucleotide variants. Whole exome sequencing of a nuclear family revealed novel single nucleotide variants of MYH7 and MKL2 in an asymptomatic father while the offspring with severe childhood-onset LVNC harbored an additional missense variant in the cardiac transcription factor, NKX2-5, inherited from an unaffected mother. Mice bred to compound heterozygosity for the orthologous missense variants in Myh7 and Mkl2 had mild cardiac pathology; the additional inheritance of the Nkx2-5 variant yielded a more severe LVNC-like phenotype in triple compound heterozygotes. RNA sequencing identified genes associated with endothelial and myocardial development that were dysregulated in hearts from triple heterozygote mice and human induced pluripotent stem cell–derived cardiomyocytes harboring the three variants, with evidence for NKX2-5’s contribution as a modifier on the molecular level. These studies demonstrate that the deployment of efficient gene editing tools can provide experimental evidence for complex inheritance of human disease.

**One sentence summary:** A combination of three inherited heterozygous missense single nucleotide variants underlying a familial congenital heart defect.

## Main Text

The genetic etiology of complex phenotypes or diseases, such as type 2 diabetes, Parkinson’s and heart disease, remain largely undetermined (*1*, *2*). High-throughput DNA sequencing has led to large databases of genetic information revealing the vast landscape of genetic variation in the absence of disease, thereby improving our ability to identify pathogenic variants involved in Mendelian disorders (*3*–*5*). However, the lack of experimental approaches to examine scenarios that involve of multiple variants and their epistatic relationships has hampered the ability to dissect the genetic mechanisms of complex phenotypes, especially those that may arise from oligo- or polygenic inheritance (*6*). Modifier genes, which likely contribute to phenotypic characteristics such as penetrance and severity, further complicate genetic association studies because of the multitude of mechanisms through which they act on molecular networks that maintain homeostasis (*7*). Congenital abnormalities represent a robust model to study complex genetic relationships. They are present at birth and, therefore, likely less dependent on environmental factors and lifestyle choices that are difficult to control for in the study of adult-onset diseases, such as coronary artery disease and type 2 diabetes.

Definitive genetic causes of congenital heart disease (CHD), the most common congenital malformation, have similarly been elusive (*8*, *9*). Rare inherited and de novo monogenic aberrations account for ~10% of cases, based on a recent exome sequencing study of CHD trios, while copy number variants have been identified in ~25% of cases (*10*, *11*). Single-gene mutations of large effect have been described through studies of relatively rare families with CHD, including mutations in transcription factors (i.e., TBX5, GATA4 and NKX2-5), signal transduction components in the NOTCH, *TGF**β*** and RAS pathways (e.g., NOTCH1, GDF11, PTEN), and structural proteins (e.g., MYH6, ACTC1) (*12*, *13*). Despite these advances, the genetic basis of the majority of CHD cases remains unknown. While multiple hypotheses have been posed to explain the paucity of de novo mutations identified in genome studies of CHD patients, such as somatic mutation or mutations in the noncoding genome, CHD has been hypothesized as a complex phenotype derived from variation in multiple genetic loci (*14*–*17*). Multiple cardiovascular defects are often identified in one individual, and phenotypes are often variable and exhibit incomplete penetrance, supporting the involvement of genetic modifiers (*14*, *15*). Mouse models of CHD support this hypothesis as genetic background can dictate the penetrance of various CHDs, and haploinsufficiency of a single gene, such as *Nkx2-5*, can lead to various phenotypes (*18*–*20*).

Recent improvements in gene editing facilitated by CRISPR-Cas proteins provide the opportunity to test hypotheses involving the potential for oligogenic inheritance of disease (*21*). In parallel, the establishment of human pluripotent stem cell models of differentiation has fostered the ability to study aberrant regulatory events that occur during embryonic development, such as those that lead to CHD, in a human context (*22*). These tools collectively create the opportunity to determine the involvement of oligogenic inheritance in CHD. Here, we utilize these advances to dissect a complex familial case of a CHD and identify a rare missense NKX2-5 variant that acts as a modifier, on both phenotypic and molecular levels, in conjunction with missense variants in the transcription factor, MKL2, and the sarcomeric protein, MYH7. To our knowledge, this work represents the first experimental validation of a human oligogenic phenotype.

### Multiple genetic mutations segregate with familial left ventricular non-compaction

Upon presentation of an infant with congestive heart failure requiring mechanical ventilation and inotropic support, echocardiography revealed severely depressed left ventricular (LV) function and deep LV trabeculations, characteristic of a disease known as left ventricular non-compaction (LVNC) (**Fig. 1A, right**). LVNC is thought to represent failure of full cardiomyocyte maturation and accounts for 2-3% of all cardiomyopathies though it has been argued its incidence is underestimated (*23*). Family history revealed a sibling who had fetal demise at 8 months of gestation due to heart failure. We obtained the pathology report and the heart, which was preserved at time of autopsy. Examination of the histologic sections revealed biventricular noncompaction, based on the deep recesses in the myocardial walls of both ventricles, right ventricular dilation, and widespread fibrosis (**Fig. 1B**). Echocardiography performed on immediate family members revealed a 4-year-old sibling who exhibited clear evidence for LVNC and cardiac dysfunction (**Fig. 1C**). The proband’s mother had normal cardiac function and anatomy, and the father, who was asymptomatic, also had normal cardiac function but some LV trabeculations that were deeper than normal and suggestive of LVNC. Extended maternal and paternal family history was negative. Collectively, these findings suggested vertical transmission of LVNC from the father with a markedly increased severity of disease in all offspring.

**Fig. 1.**
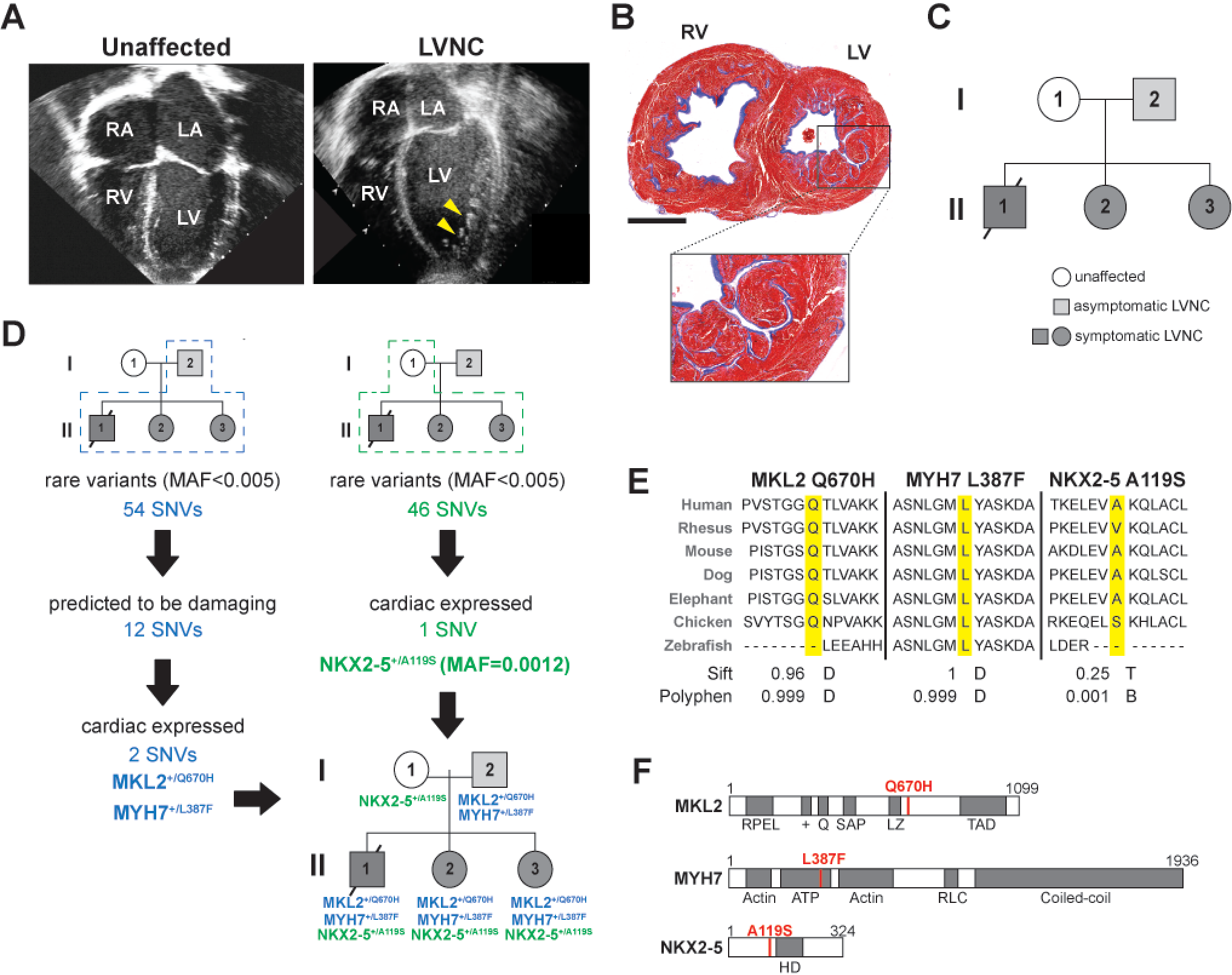
Phenotypic and Genomic Analysis of LVNC Pedigree. (**A**) Four chamber echocardiography view showing the left atria (LA), right atria (RA), left ventricle (LV) and right ventricle (RV). Unaffected at left(*66*), LVNC-affected (II-3) at right. Yellow arrows indicate trabeculations. (**B**) Transverse sections of neonatal human heart with confirmed LVNC (II-1). Scale bar = 6mm. The top image was constructed from multiple 1.5x images stitched together. (**C**) Pedigree showing inheritance pattern of LVNC. White = unaffected, light grey=asymptomatic LVNC, dark grey = symptomatic LVNC. (**D**) Schematic of exome sequencing analysis. Workflow for variants inherited from the asymptomatic father is on the left and shown in blue. Workflow for variants inherited from the unaffected mother is on the right and shown in green. Inheritance pattern of variants of interest is shown in pedigree on lower right. (**E**) Conservation of each amino acid residue is shown. The variant of interest is highlighted in yellow. Predictive effects of variants on protein function based on Sift and Polyphen programs with score of 1 being most likely to have detrimental effects. Damaging (D), tolerated (T) and benign (B).(**F**) Protein domains are illustrated for MKL2, MYH7 and NKX2-5. SNVs identified in patients are indicated in red. RPxxxEL repeat domain (RPEL), basic domain (+), glutamine rich region (Q), SAF-A, acinus, PINUS domain (SAP), leucine zipper (LZ), transcriptional activation domain (TAD), actin binding domain (actin), ATP binding domain (ATP), regulatory light chain (RLC), homeodomain (HD).

To evaluate the potential for a genetic cause of LVNC in this family, whole-exome sequencing was performed on DNA from the immediate family of the afflicted patient, including the child who suffered fetal demise. We first focused on inherited novel and rare (minor allele frequency, MAF<0.005) variants that segregated with the disease and identified 54 rare single nucleotide variants (SNVs) (**Fig. 1D, left, Table S1**). Twelve of these SNVs were predicted to be damaging, but only two were expressed in heart tissue (*24*, *25*). The first was a novel heterozygous missense variant in myosin heavy chain 7 (MYH7) involving a leucine-to-phenylalanine substitution at position 387 (L387F) (**Fig. 1D, left**). The SNV in this important sarcomeric gene was de novo in the father, as Sanger sequencing did not reveal the variant in his parents or siblings (data not shown). Variants in MYH7 are associated with LVNC, although they were generally identified through targeted sequencing of a small number of genes and often do not exhibit complete penetrance (*26*, *27*).

The second variant was a novel heterozygous missense SNV in the transcription factor *MKL2* (myocardin-related transcription factor B, MRTF-B), which is essential for vascular development (**Fig. 1D, left**) (*28*). This SNV resulted in a glutamine-to-histidine substitution at position 670 (Q670H). The myocardin family of regulators are potent transcriptional co-activators of serum response factor (SRF) and Mef2c, both essential for cardiogenesis (*29*). *MKL2* deletion leads to defects in the cardiovascular epithelial-to-mesenchymal transition during embryonic development and cancer metastasis through interaction with the YAP-TEAD and *TGF**β*** pathways (*30*, *31*). While it has not previously been associated with cardiac disease, variants that presumably cause loss of function are not tolerated based on data analyzed by the Exome Aggregation Consortium (pLI = 0.99) (*4*). Conversely, its homolog *MKL1,* whose redundant function is often assessed simultaneously with *MKL2*, does not exhibit a similar level of intolerance to loss of function (pLI = *0.54*) (*32*). Luciferase assays using an SRF-dependent *TGF**β**2* promoter revealed MKL2 Q670H had partially reduced transcriptional activity compared to Wt, providing evidence for dysfunction induced by this variant (**Fig. S1**).

Given the marked increase in severity of disease in the three children, we also posed a hypothesis aimed at identifying variants inherited from the unaffected mother that might serve as genetic modifiers of the LVNC phenotype (**Fig. 1D, right**). Inclusion of rare and common variants inherited by all three affected children from the unaffected mother resulted in identification of 47 SNVs (**Table S1**). Filtering using expression data and conservation discerned a rare heterozygous missense variant in *NKX2-5*, resulting in an alanine-to-serine substitution at position 119 (A119S) (*24*). This variant has a comprehensive MAF of 0.0012 (**Fig. 1D, right**), but is twice as common in South Asians (MAF=0.0026). NKX2-5 is one of the “core” transcriptional regulators of cardiac development and heterozygous mutations in the DNA-binding homeodomain have been associated with CHD previously, including LVNC (*33*, *34*). The A119S variant is outside the homeodomain, but was reported to cause limited DNA-binding in vitro and has been noted in the setting of CHD, although with incomplete penetrance (*35*–*37*). The variant in NKX2-5, along with MYH7 and MKL2, were validated by Sanger sequencing (**Fig. S2, Table S2**).

MYH7 and MKL2 SNVs affect highly conserved amino acids, suggesting significant evolutionary pressure on these residues (**Fig. 1E**). The NKX2-5 SNV is not conserved on an amino acid level in Rhesus macaque, though the lack of polarity at this site is maintained by the amino acid substitution of alanine to valine (**Fig. 1E, top**); nevertheless, the reduced conservation and the 0.1% allele frequency suggest tolerance of variability at this residue. Prediction of functional damage to the protein induced by each variant suggests that both MKL2 and MYH7 SNVs are likely damaging, while the NKX2-5 SNV is predicted to be tolerated, in agreement with its conservation (**Fig. 1E, bottom**). MYH7 L387F is located within the critical ATP binding domain, while the SNVs in MKL2 and NKX2-5 are not within known protein domains (**Fig. 1F**). Subsequent sequencing of all paternal family members revealed the presence of the MKL2 Q670H variant in an unaffected uncle of the proband and grandfather, indicating that this variant is not sufficient to cause LVNC (**Fig. S2B**). The observation of isolated MKL2 Q670H or NKX2-5 A119S variants in unaffected individuals raised the possibility that these alterations have subtle effects on protein function and act as modifiers of the phenotype in the presence of the MYH7 L387F variants (*38*). While the inheritance pattern of the heterozygous missense variants in MKL2, MYH7 and NKX2-5 is consistent with an oligogenic contribution to the severe disease phenotype, we sought to experimentally test this hypothesis in vivo in mice and in vitro in human induced pluripotent stem cell (hiPSC)–derived cells.

### Functional significance of MYH7, MKL2, and NKX2-5 missense variants in vivo

To evaluate the functional consequence of the MKL2, MYH7 and NKX2-5 amino acid substitutions identified in the LVNC family, we generated mice (C57BL/6J) harboring the orthologous variants by CRISPR-Cas9 (**Fig. 2A-C**). Mice heterozygous for each allele individually were normal physiologically. Based on RNA sequencing of heterozygous mouse hearts, targeted loci did not exhibit changes in mRNA splicing of the targeted genes or a major allelic imbalance (**Fig. S3**). Animals were then bred to homozygosity to test the functional consequence of each missense variant. Although Myh7^L387F^ heterozygous animals were observed at the expected mendelian ratio, no homozygous mutant animals were detected in offspring from heterozygous intercrosses. By sacrificing neonates and embryos at progressive stages of development, we found that homozygous animals were embryonic lethal by embryonic day (e) 10.0 with large pericardial edema, indicative of heart failure (**Fig. 2D**). These observations demonstrated that the MYH7 L387F substitution is damaging, as predicted.

**Fig. 2.**
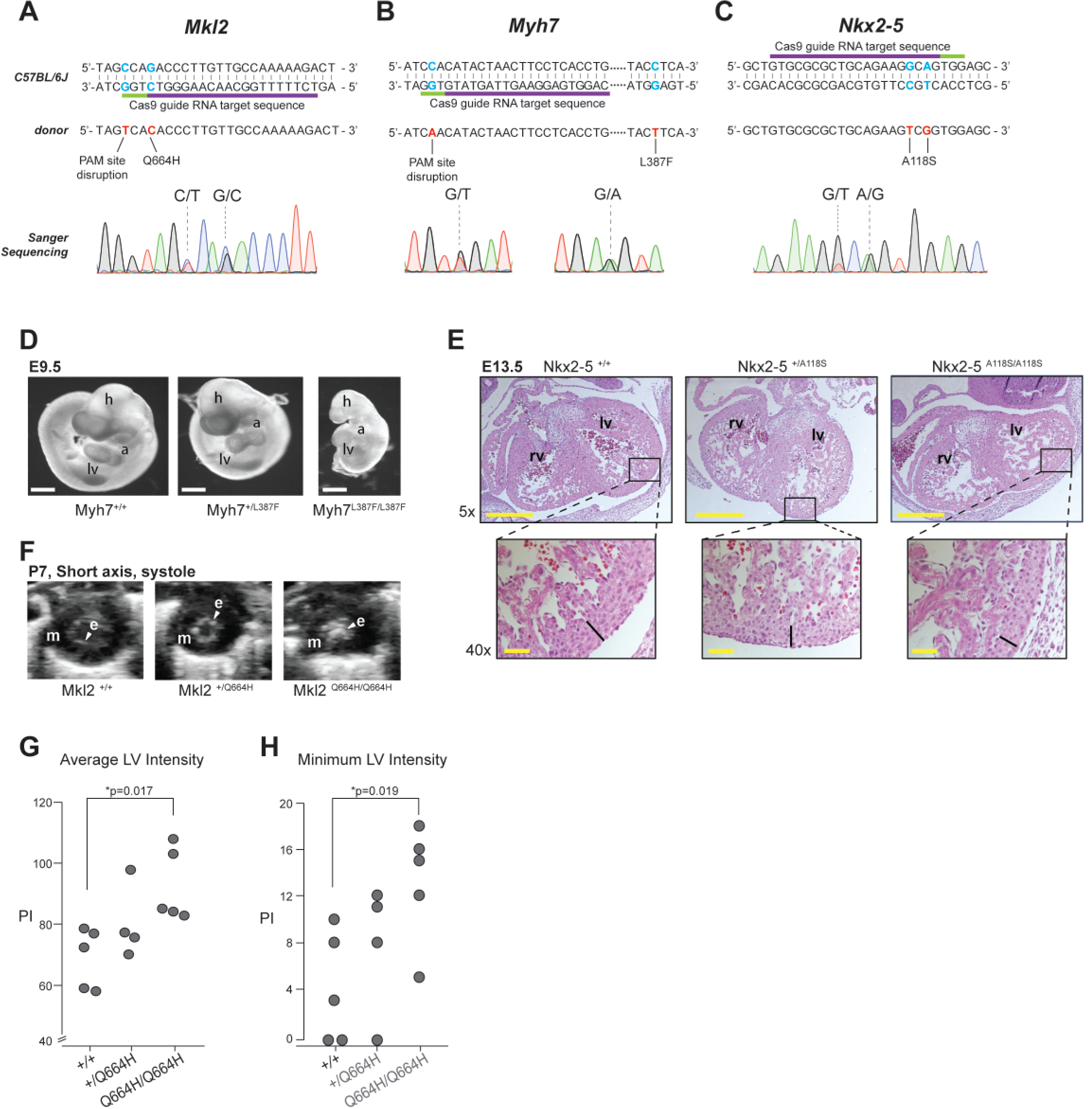
In Vivo Evaluation for Functional Consequences of Single Nucleotide Variants Segregating with Disease in LVNC Pedigree. (**A-C**) Cas9 targeting scheme is shown for each orthologous locus in the mouse. The genomic sequence is at the top. Cas9 targeting sequence is indicated in purple. PAM site is in green. Blue nucleotides are mutated in targeted mice. Single-stranded donor sequence is in the middle bottom with mutated nucleotides indicated in red. Sanger sequencing traces at each locus are shown for heterozygous mice at bottom. Guide RNA sequences and donor templates are in **Table 2**. (**D**) Left lateral images of E10.0 embryos that result from a Myh7^+/L387F^ × Myh7^+/L387F^ cross are shown with genotypes labeled at bottom. Growth failure and enlargement of heart are evident in the homozygous mutant. h, head; lv, left ventricle; a, atrium. Scale bar = 5mm. **(E) Transverse histologic sections** of E13.5 embryos with indicated genotypes. Width of compact layer indicated by black bar. RV, right ventricle; LV, left ventricle. 5x at top, scale=500um. 40x at bottom, scale =50um. (F) Short axis view of left ventricle in systole viewed by echocardiography of P7 mice with indicated Mkl2 genotypes. m, myocardium signified by dark area; e, endocardium (white area) indicated by arrowheads. (**G**) Average pixel intensity (PI) within the left ventricle was calculated from three litters of mice. P-value was calculated using a T-test. Average pixel intensity (PI) within the left ventricle of a P4 mouse heart calculated from three independent litters. (**H**) Minimum PI was calculated as in G.

Mendelian ratios were observed for Nkx2-5^A118S^ or Mkl2^Q664H^ heterozygous crosses, and homozygous animals did not exhibit evidence of cardiac failure. However, homozygotes for either residue exhibited abnormalities in the ventricular wall before the first week of life. At embryonic time points, we noted a thin left ventricular myocardial wall in Nkx2-5^A118S^ homozygous mice as well as thicker trabeculae (**Fig. 2E**). These abnormalities were subtle and resolved over time as apparent differences were not observed postnatally. While there was no indication of cardiac dysfunction by echocardiography, we noted marked echogenic foci in the left ventricular cavity of postnatal day 3 (P3) Mkl2^Q664H^ homozygous mice similar to that in human patients with LVNC (**Fig. 2F**). Quantification of average pixel intensity, as well as the minimum intensity within the entire LV area, confirmed greater levels of signal in heterozygous and homozygous animals than Wt, indicative of increased endocardial surface typically observed in the setting of deep ventricular trabeculations (p=0.017, p=0.019, T-test, **Fig. 2G, H**). Interestingly, this phenotype also resolved with age as echogenic foci were not present by 2 weeks of age (data not shown). Cardiomyocyte-specific deletion of Mkl2 did not reveal a similar phenotype, suggesting it is essential for proper development of another cell type within the heart (*40*). These results indicate that the MKL2 and NKX2-5 SNVs in patients adversely affect the ability of the encoded protein to promote timely ventricular development in a mouse model, although both exhibit milder effects than the MYH7 variant.

### Triple compound heterozygote mice exhibit LVNC

Having established that each of the Myh7, Mkl2 and Nkx2-5 missense variants was consequential in vivo upon homozygosity, we intercrossed heterozygous mice to determine if compound heterozygosity of the three genetic mutations could cause an LVNC-like phenotype similar to that observed in humans. Mild hypertrabeculation and some subtle apical recesses were observed in Myh7^L387F/+^Mkl2^Q653H/+^ mice (**Fig. 3A**). Remarkably, Myh7^L387F/+^Mkl2^Q664H/+^Nkx2-5^A119S/+^ mice had clear abnormalities upon pathologic examination. Immunohistochemistry with an antibody to the endocardial marker, endomucin, at P3 revealed deep trabeculations in the left ventricle wall very similar to that seen in patients with LVNC and as observed in the autopsy of the affected child in the family described here (**Fig. 3A, left**). Deep recesses in the apex of the left ventricle were also observed in the triple mutants at P3 (**Fig. 3A, center**). As often seen in humans, the left ventricular wall continued to develop and by P7 the myocardial walls had further compacted, and the increased trabeculation had resolved in Myh7^L387F/+^Mkl2^Q664H/+^ and Nkx2-5^A118S/+^ mice. In contrast, the hypertrabeculation and apical recesses persisted in triple heterozygous mice, consistent with Nkx2-5 functioning as a genetic modifier of Myh7 and Mkl2 effects on ventricular development (**Fig. 3A, right**).

**Fig. 3.**
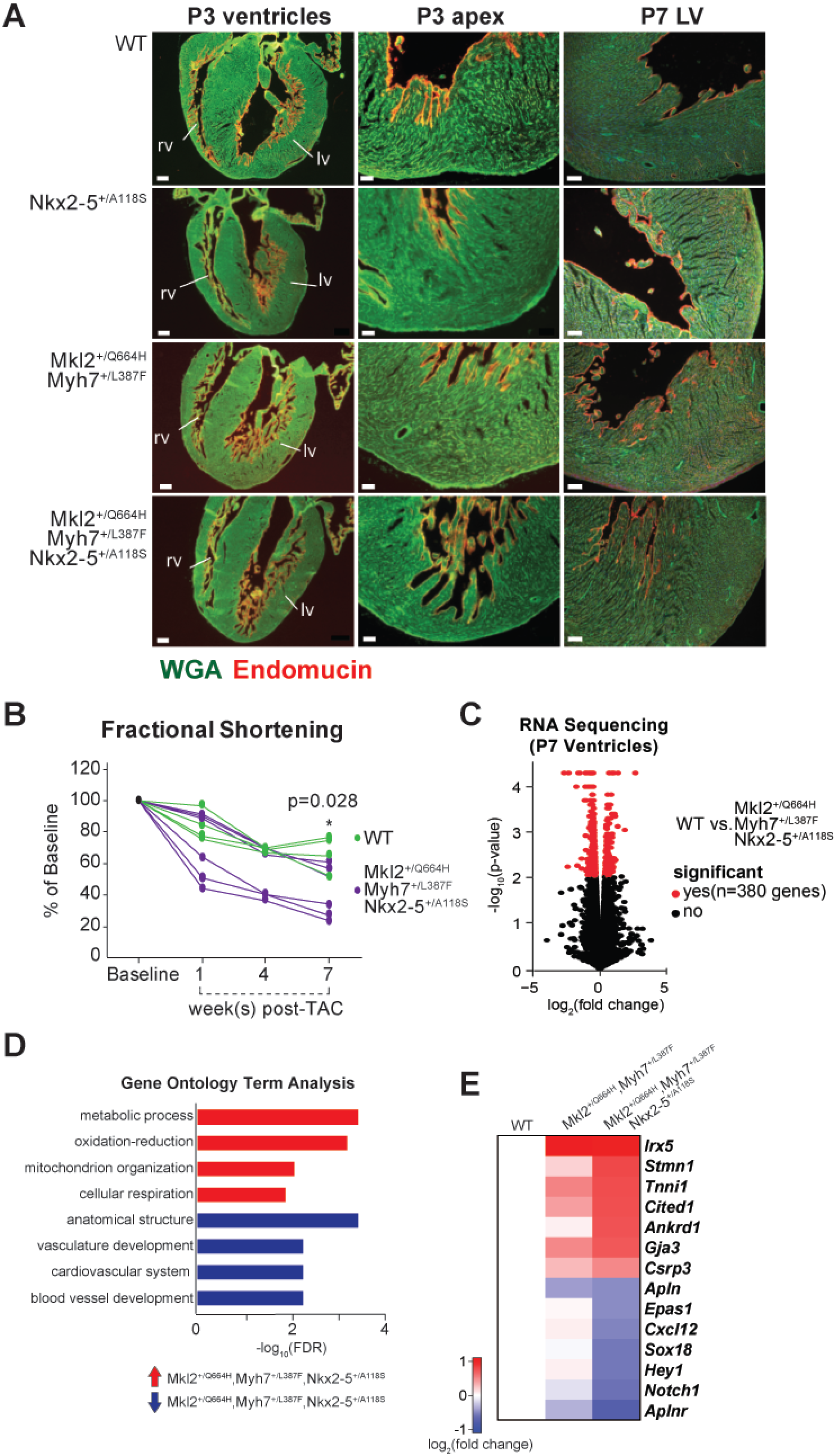
Mkl2:Myh7:Nkx2-5 Compound Heterozygous Mice Recapitulate LVNC on a Phenotypic and Molecular Level. (**A**) Left: right ventricle (rv) and left ventricle (lv) observed in four chambered views of P3 mice at 2.5x. Scale bar = 200 mM. Center: the apical region of the same hearts shown on left imaged at 10x Scale bar = 50 mM. Right: Left ventricle of P7 mice imaged at 10x. Scale bar = 200 mM. (B) Fractional shortening of wild-type (Wt) or Mkl2^+/Q664H^ Myh7^+/L387F^ Nkx2-5^+/A118S^ mice exhibited as percent of baseline before and after transverse aortic constriction (TAC). (**C**) Volcano plot for differential gene expression analysis between Wt and triple heterozygous mouse P7 ventricles. Statistical significance is indicated in red (n = 380 genes). (**D**) Gene Ontology term analysis of differentially expressed genes. Barplot depicts −log_10_(FDR). Red indicates genes detected at higher levels in triple compound heterozygous animals, and blue indicates categories of genes that are expressed at lower levels in triple compound heterozygous animals, compared to Wt, by a fold change of > 0.3. Full list can be found in **Table S3**. (**E**) Heatmap of selected genes depicting fold change of FPKM between Wt and Mkl2^+/Q664H^ Myh7^+/L387F^ double compound heterozygous mice or Wt and Mkl2^+/Q664H^ Myh7^+/L387F^ Nkx2-5^+/A118S^ triple compound heterozygous mice. Key at lower left indicates log_2_ fold change.

Echocardiography did not detect cardiac dysfunction at baseline in adult Myh7^L387F/+^Mkl2^Q664H/+^Nkx2-5^A118S/+^ mice. Given reports that LVNC can be exacerbated under stress only to resolve when the external condition, such as pregnancy or intense physical exertion, is removed, we tested whether physiological stress would induce cardiac dysfunction in triple heterozygous mice (*41*, *42*). Transverse aortic constriction (TAC) on triple heterozygous or Wt mice was used to increase the pressure load on the left ventricle, creating hemodynamic stress (*43*). Triple heterozygous mice exhibited statistically significant reductions in the fraction of blood ejected from the LV with each beat (ejection fraction) and the degree of shortening of the LV diameter with each contraction (shortening fraction), compared to Wt mice (p=0.028, **Fig. 3B**). This pathological response was 60–70% penetrant, suggesting that the triple heterozygous mice were on the threshold of functional abnormality, despite the fully penetrant morphologic abnormality of LVNC.

We next sought to understand the molecular changes induced by the Myh7^L387F/+^Mkl2^Q664H/+^ and Myh7^L387F/+^Mkl2^Q664H/+^Nkx2-5^A118S/+^ variants in the mouse hearts. RNA sequencing of tissue from the apex of P7 hearts revealed subtle yet statistically significant upregulation of genes associated with metabolism in triple compound heterozygous mice, whereas genes associated with vasculogenesis were downregulated in these samples based on gene ontology analysis (GO terms) (**Fig. 3C, D, Table S3,** p-value < 0.005). Increased expression of genes involved in mitosis was also observed, such as *Stmn1*, though quantification of phospho-histone 3 foci at postnatal timepoints did not result in a statistically significant difference between genotypes, suggesting productive cell division is not increased in triple compound heterozygous mice (data not shown). Genes expressed at higher levels in the myocardial trabeculae during embryonic development, such as *Irx5,* were also upregulated in triple mutant mice, supporting the histological observation of hypertrabeculation (**Fig. 3G, Table S4**). Genes associated with earlier stages of development and directly regulated by Nkx2-5, such as *Ankrd1,* were additionally expressed at higher levels in triple compound heterozygous mice providing molecular evidence of their less mature state compared to wild-type (*44*, *45*). Conversely, genes associated with endothelial cell development, such as *Hey1*, *Notch1*, and *Sox18,* were downregulated (**Fig. 3G, Table S4**). Previous work suggested that disruption of Notch signaling during embryonic development contributes to improper endothelial cell development and subsequent LVNC through reciprocal interactions with the myocardium, in agreement with a model in which a combination of heterozygous SNVs contributes to the disease phenotype (*46*). Moreover, the low expression of genes necessary for mature vascular endothelial cell function in the triple heterozygous animal suggests poor endothelial cell function is associated with the LVNC phenotype in this mouse model. RNA sequencing after cardiomyocyte-specific deletion of Mkl1/2 hearts similarly revealed dysregulation of epithelial cell-related pathways, further supporting our results (*40*).

### Human iPSC-derived cardiomyocytes exhibit disease pathology

To determine if the phenotype exhibited by triple heterozygous mice reflected the effects of the genetic mutations in human cardiomyocytes, we generated patient-specific human pluripotent stem cells (hiPSCs) from multiple family members and differentiated them to cardiomyocytes via WNT pathway modulation (*47*). Differentiation efficiency based on cardiac troponin T expression was similar in control and patient lines; however, discrepancies in adherence between patient lines became apparent by day 7 of differentiation when cardiomyocytes began to beat (**Fig. 4 A, B**). Cells derived from an individual with symptomatic LVNC (MYH7^L387F/+^MKL2^Q670H/+^NKX2-5^A119S/+^) formed aggregates that were loosely attached to the cell-culture dish (**Fig. 4B**). This phenomenon was less apparent in the asymptomatic case (MYH7^L387F/+^MKL2^Q670H/+^) and undetectable in the unaffected case (NKX2-5^A119S/+^), correlating with the presence of all three SNVs studied here. Previous work concluded Mkl2 positively regulates expression of genes necessary for adhesion, in agreement with our observation in hiPSC-derived cardiomyocytes (*30*).

**Fig. 4.**
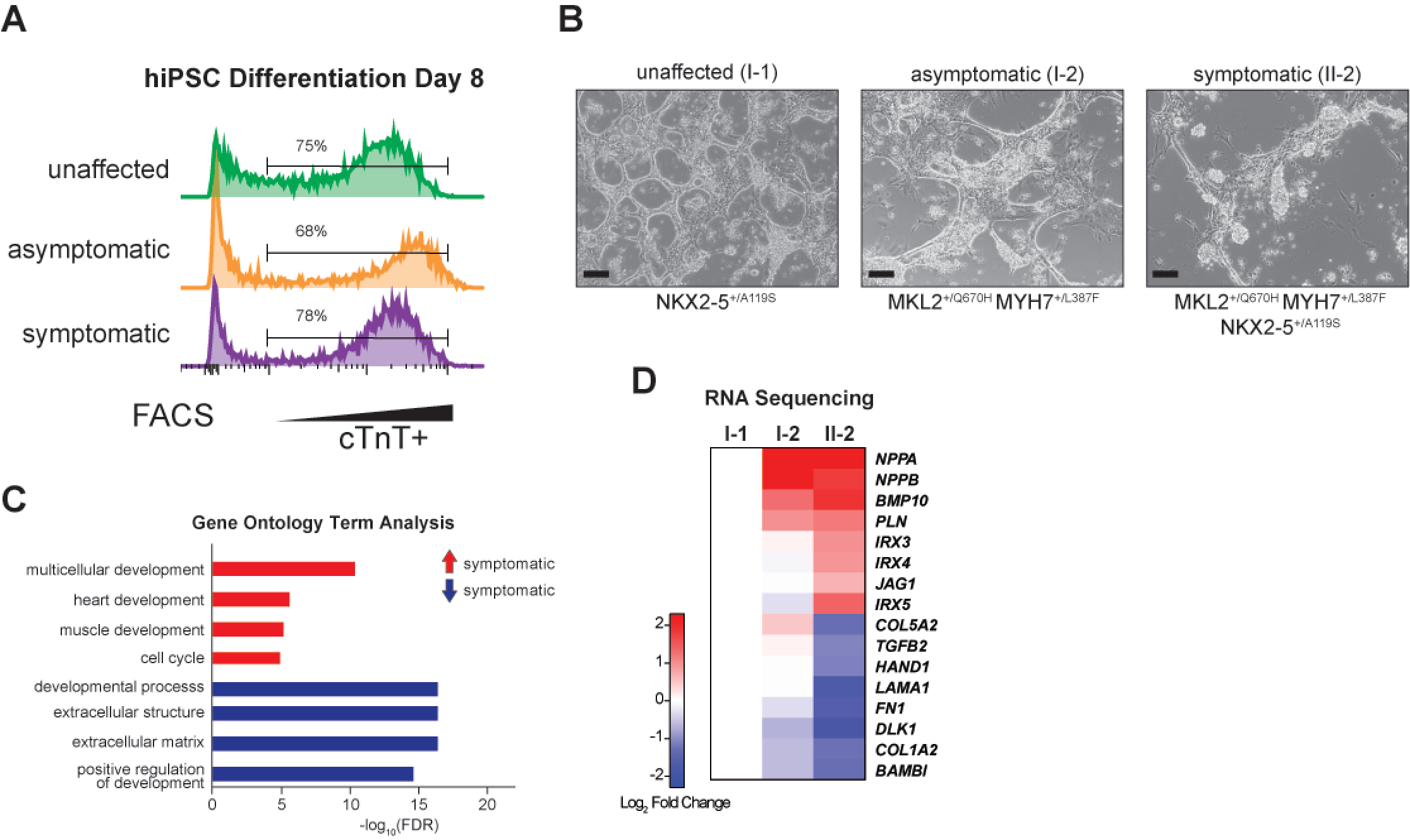
hiPSC Differentiation towards Cardiomyocytes Replicates Dysregulation Identified in Triple Compound Heterozygous Mice. (**A**) FACS analysis of hiPSC-derived cells collected at day 8 after induction of cardiac differentiation and stained for cardiac troponin T (cTnT). (**B**) Brightfield images of hiPSC-derived cardiomyocytes from three family members as indicated. Scale = 100 mM. (**C**) Gene Ontology term analysis of differentially expressed genes. Barplot depicts −log_10_(FDR). Red indicates genes more highly expressed in cells from the symptomatic individual (Mkl2^+/Q664H^ Myh7^+/L387F^ Nkx2-5^+/A118S^) compared to cell from the unaffected (Nkx2-5^+/A118S^), and blue indicates categories of genes expressed at lower levels in the symptomatic individual, by a fold chance of > 0.3. Full list can be found in **Table S5**. (**D**) Heatmap depicting log2 fold change of asymptomatic (I-2) and symptomatic (II-2) LVNC hiPSC-derived cardiomyocytes, compared to unaffected (I-1). Key at lower left. Full list can be found in **Table S6**.

To understand the transcriptional landscape of human cardiomyocytes from family members, RNA sequencing was completed at day 8 of differentiation towards cardiomyocytes. Gene ontology (GO) term analysis revealed dysregulation of genes sets involving upregulation of cell cycle and mitosis in cells derived from the symptomatic LVNC case (MYH7^L387F/+^MKL2^Q670H/+^NKX2-5^A119S/+^), similar to that observed in triple heterozygous mice (**Fig. 4C, Table S5,** FDR adjusted p-value < 0.05). Interestingly, GO term analysis on genes that were downregulated revealed categories associated with cell adhesion and extracellular matrix deposition, supporting the visual observation of a lack of adhesion in the early-onset case (**Fig. 4B**). Evaluation of genes associated with the cardiac progenitor state, such as *BMP10, IRX3, IRX5* and *NPPA*, exhibited higher expression in the LVNC lines than the unaffected line (**Fig. 4D, Table S6**). Conversely, genes associated with adhesion (e.g., *FN1, COL1A1, COL5A1*) showed lower expression in the symptomatic cell line than the unaffected (NKX2-5^A118S/+^) (**Fig. 4D**, p<0.005). In agreement with the mouse transcriptional data that included the Nkx2-5 A118S SNV, the fold changes of genes upregulated and downregulated, compared to unaffected, were greater in the cell line derived from the individual diagnosed with early onset LVNC and harboring all three genetic variants (**Fig. 4D**). Thus, the transcriptional landscape of the LVNC phenotype could be recapitulated through directed differentiation of hiPSCs, and the modifying effect of the NKX2-5 variant on the disease phenotype was confirmed on a molecular level. Additionally, these results suggest that, while disruption of proper endothelial cell development and function may contribute to LVNC, there is a cardiomyocyte cell-autonomous component given that endothelial cells are not present in our differentiations.

## Discussion

The contribution of genetic modifiers to disease has traditionally been difficult to prove experimentally. Here, we present the first evidence of a single nucleotide variant that modifies a congenital heart disease caused by multiple variants. CRISPR-Cas9 was used to generate mice encoding three SNVs orthologous to those identified in LVNC patients, and compound triple heterozygotes recapitulated the disease histologically. Transcriptome analysis of afflicted mice further revealed dysregulated pathways previously associated with ventricular maturation through traditional mouse genetic studies. Importantly, the Nkx2-5 SNV A118S exacerbated this dysregulation, supporting the conclusion that this SNV acts as a modifier in the case of LVNC caused by missense variants in MYH7 and MKL2. In agreement with the mouse data, differentiation of hiPSCs towards cardiomyocytes derived from the patient with symptomatic LVNC or all three mutations revealed persistent expression of genes associated with the immature trabecular myocardium. Our data collectively suggest this case of LVNC is inherited in an oligogenic fashion, providing the first experimental validation of a CHD inherited in this manner, to our knowledge.

The development of assays to test variants of unknown significance is essential for the advancement of precision medicine initiatives (*48*, *49*). While deletions, insertions and frameshift variants have somewhat predictable consequences, the effect of millions of single nucleotide variants identified in each individual’s genome are difficult to assess computationally (*50*–*52*). Recent advances in gene editing technologies have now created an avenue to interrogate the contribution of these variants to phenotypes and disease (*53*). By creating a mouse model encoding orthologous mutations, our data show that traditional metrics for the identification of phenotype-associated variants are likely not designed to identify the subtle effects of genetic modifiers because they rely heavily on sequence conservation and structural information, which is not always available for genes of interest (*54*). This result argues experimental exploration and validation are currently an essential step in the search for modifiers of disease (*48*, *49*). Accordingly, additional experimentation and analysis must be done to understand the prevalence and context in which NKX2-5 A119S can act as a modifier of CHD. Multiple relatively rare missense variants in transcription factors and signaling pathway modulators essential for cardiovascular development have been identified in the population, but their involvement in the approximately 65% of unresolved CHD cases has not been widely appreciated (*4*). Similarly, missense variants in MYH7, sometimes even affecting neighboring residues, have been associated with a wide spectrum of cardiomyopathies, including hypertrophic and dilated cardiomyopathies, as well as LVNC, highlighting another scenario that may involve genetic modifiers influencing the consequences of MYH7 mutations (*55*).

Additionally, using human genetic variation to dissect biological processes will undoubtedly reveal novel insights regarding disease mechanisms. Our data suggest that while LVNC manifests itself in the myocardium, it can also be associated with improper development of the endothelium. More screening needs to be done to understand whether the putative mechanism involved in this case of LVNC is unique and should be considered an “n of 1,” or if dysregulation of endothelial cell biology is a common factor in disease pathogenesis (*56*). As we refine our understanding of the regulatory mechanisms that govern cell-autonomous and non-cell-autonomous cellular states using advances in both gene editing and single cell next-generation sequencing approaches, our ability to correlate genetic variation with phenotypic outcome will improve, bringing precision medicine closer to reality.

## Acknowledgements

The authors thank the Srivastava lab and members of the Gladstone Institute of Cardiovascular Disease for helpful discussion and feedback, and the Gladstone Histology and Light Microscopy Core, Gladstone Stem Cell Core, Gladstone Transgenic Core for making this work possible. Importantly, we’d like to thank the family who generously offered to be part of our study. C.A.G. is a HHMI fellow of the Damon Runyon Cancer Research Foundation (DRG-2206-14). D.S. is supported by NHLBI/NIH grants (R01 HL057181, U01 HL098179, U01 HL100406) the Roddenberry Foundation, the L.K. Whittier Foundation and the Younger Family Fund. This work was also supported by NIH/NCRR grant C06 RR018928 to the Gladstone Institutes. All data will be available on GEO upon publication.

## Contributions

C.A.G. and D.S. conceived the study, interpreted the data and wrote the manuscript. C.A.G. and S.K.W. analyzed exome sequencing data. C.A.G., S.R., Y.D.S., and H.S., collected embryos, sectioned tissue and stained slides. C.A.G. and R.S. performed genotyping of mice. C.A.G and R.S. cultured and differentiated hiPSCs. K.M. and Y.B. collected tissue for exomes and generated hiPSCs. C.A.G. and P.Z. cloned vectors and performed luciferase assays. Y.Z. and H.S. performed echocardiography and analyzed data. A.E. imaged heart sections. P.U. examined histological sections from human heart.

## Supplementary Materials

### Materials and Methods

#### Processing of Patient Samples

DNA was extracted from venous blood using the Qiagen DNA Mini kit. DNA was extracted from the formalin-fixed paraffin embedded block using the RecoverAll Total Nucleic Acid Isolation Kit for FFPE (Thermo Fisher, 1975). DNA was submitted to either Complete Genomics (San Jose, CA) or Centrillion Technologies (Palo Alto, CA) for exome library preparation using the Agilent SureSelect Human Exon V4 enrichment kit. All libraries were sequenced to an average depth of 100X. Variants of interest were confirmed using traditional PCR amplification and Sanger sequencing. See Table 2 for primers. The FFPE-block containing the human heart was imaged with a Versa Slide Scanner at 1.5x.

#### Exome Analysis

Reads were aligned to hg19 using BWA and data was analyzed using the GATK Best Practices and Workflows (*57*). Variants were annotated using ANNOVAR, dbSNP (v138), 1000 Genomes (October, 2014) and Encode (*58*). Variants with an observed MAF < 0.005 were evaluated.

#### Luciferase Assay

Genes were cloned into plasmid pFC27A (Promega, #G8421). The *TGF**β**2* promoter was cloned into a NanoLuc genetic reporter plasmid (Promega, #N1001). Plasmids were transfected into COS1 cells using Fugene HD (Promega, #E2311). A plasmid constitutively expressing firefly luciferase was co-transfected to estimate the transfection efficiency of all plasmids. Primers for the construction of the MKL2 and *TGF**β**2* plasmids can be found in **Table s2**. The c-terminus truncation of MKL2 was generated based on alignment with MYOCD. Amino acids 826 through 1099 were deleted.

#### Mouse Embryo Targeting and Line Generation

The Nkx2-5 A118S and Mkl2 Q664H mice were generated by the Jackson Laboratory CRISPR-Cas9 mouse model generation service while the Myh7 L387F mouse was generated by the Gladstone Transgenic Core. Each line was made in the C57BL/6J background using a
200 base pair (bp) single stranded unmodified oligonucleotide (oligo) donor. In both scenarios, Cas9 protein, in vitro transcribed guide RNA and the single stranded oligo donors were microinjected at the 1 cell stage. Prior to microinjection, the efficiency of three putative guide RNA sequences was tested in mouse embryonic stem cells and evaluated using the TIDE analysis software (*59*). The most efficient guide was chosen for microinjection. Synonymous mutations were included in each donor to ensure Cas9 did not cut the donor or the locus after homologous recombination occurred. After founders were born, they were backcrossed for three generations before phenotypes were evaluated. See **Table 2** for primers used in genotyping.

#### Mouse Studies

To examine hearts at prenatal timepoints, entire embryos were fixed overnight at 4°C in 4% paraformaldehyde. They were then washed twice in PBS and stored in 70% ethanol until paraffin embedding. For postnatal timepoints, hearts were removed from the embryo and fixed overnight at 4°C in 4% paraformaldehyde, washed in PBS for 30 minutes at room temperature, incubated in 30% sucrose for 18 hours at 4°C, flash frozen in OCT (Fisher Healthcare, #23730-571) and stored at −80°C until sectioning.

For immunohistochemistry, two heart sections were placed on Superfrost Plus Microscope slides then stained with Wheat Germ Agglutinin Alexa Fluor 488 (Thermo Fisher, #W11261) and Endomucin (Abcam, #ab106100). Imaging was done on a Zeiss A1 microscope.

Echocardiography and transverse aortic constriction were conducted as previously described (*60*). The intensity of echogenic foci was quantified using ImageJ.

#### hiPSC Line Generation, Characterization and Differentiation

Patient-derived fibroblasts were cultured and reprogramming using episomal plasmids encoding OCT3/4, SOX2 and KLF4 as previously described (*61*). Cytogenic analysis confirmed normal karyotypes of all derived hiPSC lines. Pluripotency was confirmed based on staining for the transcription factors NANOG (Abcam, #ab21624), OCT3/4 (Abcam, #ab19857) and SOX2 (Abcam, #ab97959), as well as the surface markers TRA-1-60 (Millipore, #MAB4360) and SSEA4 (Abcam, #ab16287). hiPSCs were maintained in feeder free culture on Corning Matrigel hESC-Qualified matrigel (BD, #354277) using Essential 8 Medium (Thermo Fisher, #A1517001). Passaging was done with accutase (Stem Cell Technologies, #07920) and Y-27632 (ROCK inhibitor, Stem Cell Technologies, #72302) was included for 24 hours after dissociation.

To induce differentiation towards cardiomyocytes, the WNT modulation protocol was utilized with slight modifications. The WNT agonist, CHIR99021 (Stemgent Incorporated, #04-0004), was added for the first 48 hours at 6uM in RPMI (Thermo Fisher, #21870-076) with B27 supplement minus insulin (Thermo Fisher, #A1895601). Then, the WNT antagonist IWP4 (Stemgent Inc, #04-0036) was added at 5uM final concentration in RPMI with B27 supplement minus insulin for 48 hours. Cells were dissociated on day 5, and replated on plates coated with 12.5ug/mL fibronectin diluted in PBS (Sigma, #F1141). ROCK inhibitor was included for 24 hours after dissociation. Cells were subsequently cultured in RPMI with B27 supplement minus insulin until day 10 when they were switched to RPMI with B27 supplement plus insulin.

#### RNA Extraction, Sequencing Library Construction and Analysis

Mouse heart tissue was first dissociated using the Next Advance Bullet Blender. RNA was extracted from dissociated mouse heart tissue and hiPSC-derived cells using the Direct-zol kit (Zymo, #R2051) and treated with DNaseI. Each RNA sequencing experiment included three biological replicates.

RNA sequencing libraries were constructed using the 500ng of RNA and the SMARTer Stranded Total RNA Sample prep kit (Takara, #634876). Ribosomal RNA was depleted using RiboGone Mammalian (Takara, #634847). Libraries were pooled with a final concentration of 10% PhiX included (Illumina, FC-110-3001) and 100bp paired-end sequencing was obtained on the HiSeq 4000. All libraries were sequenced to a read depth of at least 2×10^7^ aligned reads per sample. Reads were aligned with STAR to hg19 or mm10 using the Ensemble iGenome gene annotation (*62*). The Tuxedo tools were used for transcript quantification and differential expression analysis and genes were quantified using fragments per kilobase per million reads mapped (FPKM) (*63*). A raw p-value < 0.005 was used to identify genes for downstream analysis in the mouse experiments while a FDR adjusted p-value (q-value) < 0.05 was used for hiPSC experiments. GO term analysis was done using the Panther Database and significance was assessed based on the FDR calculated using the Fisher’s exact test (*64*, *65*).

**Fig. S1.**
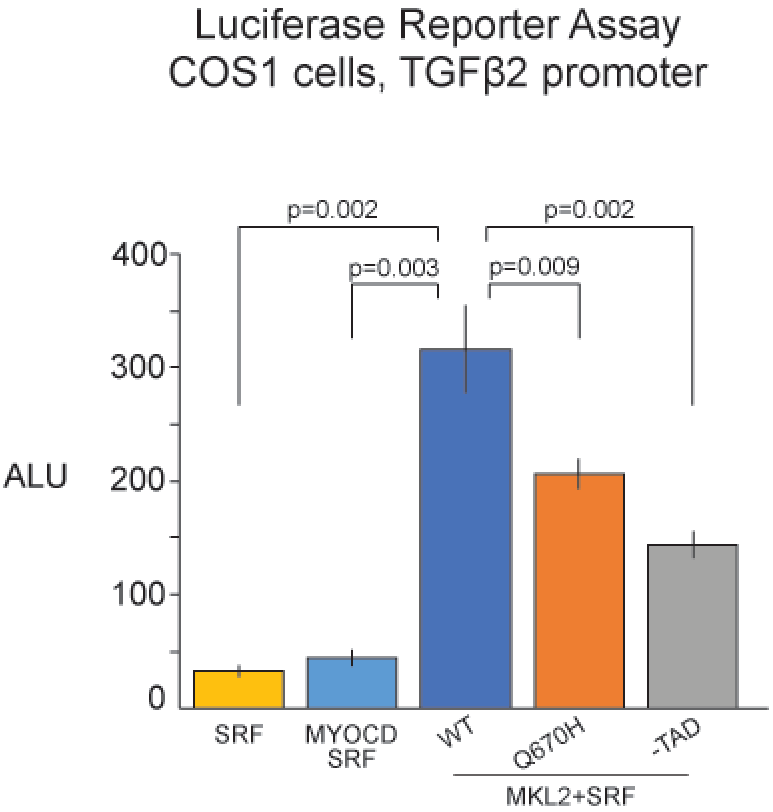
Luciferase assay reveals hypomorphic transcriptional activity of MKL2 Q670H. Outcome of luciferase assay in COS1 cells using the *TGF**β**2* promoter to measure transcriptional activity of MKL2. Normalized artificial luciferase units (ALU) are shown on the Y-axis. MYCOD and a plasmid encoding MKL2 with deletion of the transcriptional activation domain (-TAD) were included as controls. SRF was included in all experiments.

**Fig. S2.**
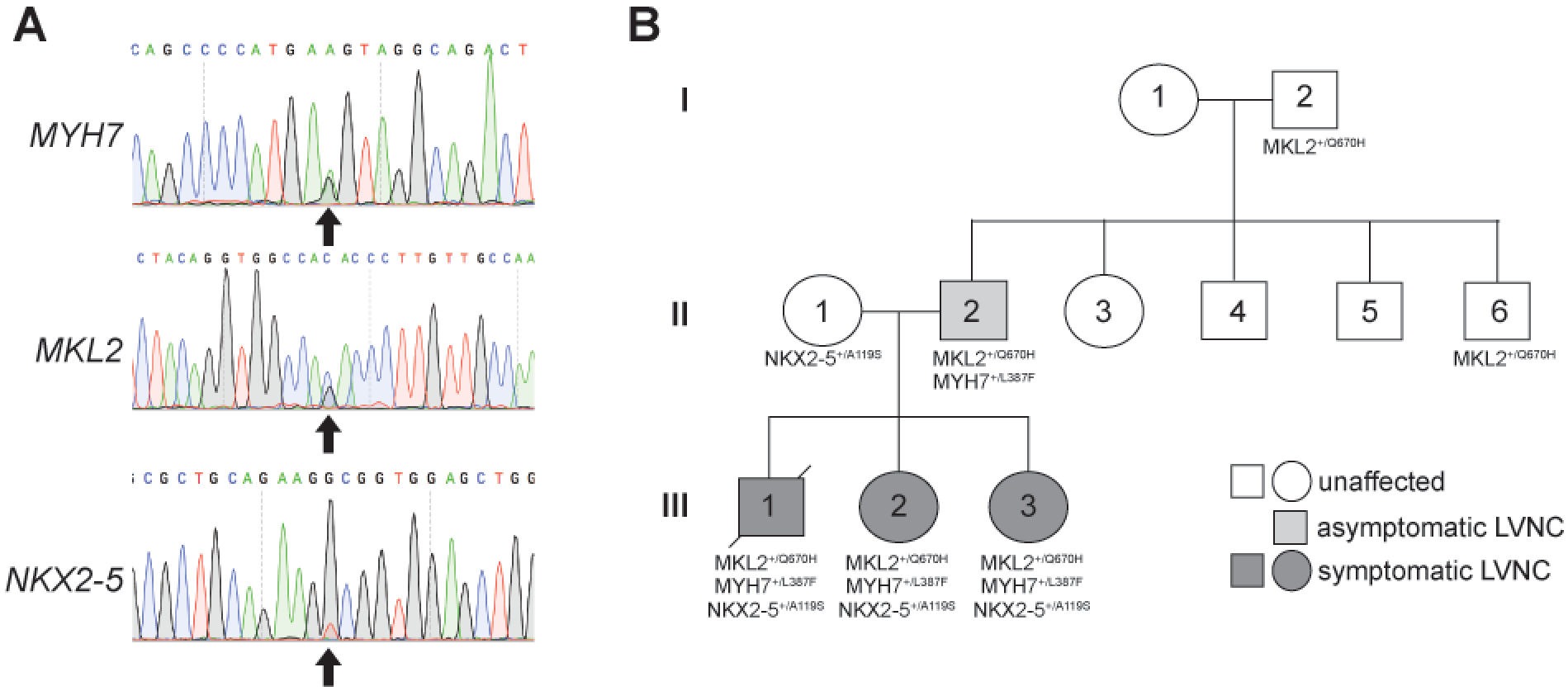
Sanger sequencing and inheritance pattern of variants of interest. **(A)** Sanger sequencing traces are shown for the three variants of interest. Variants are indicated by black arrows. Primers can be found in Table S2. **(B)** Extended family tree with identified genotypes.

**Fig. S3.**
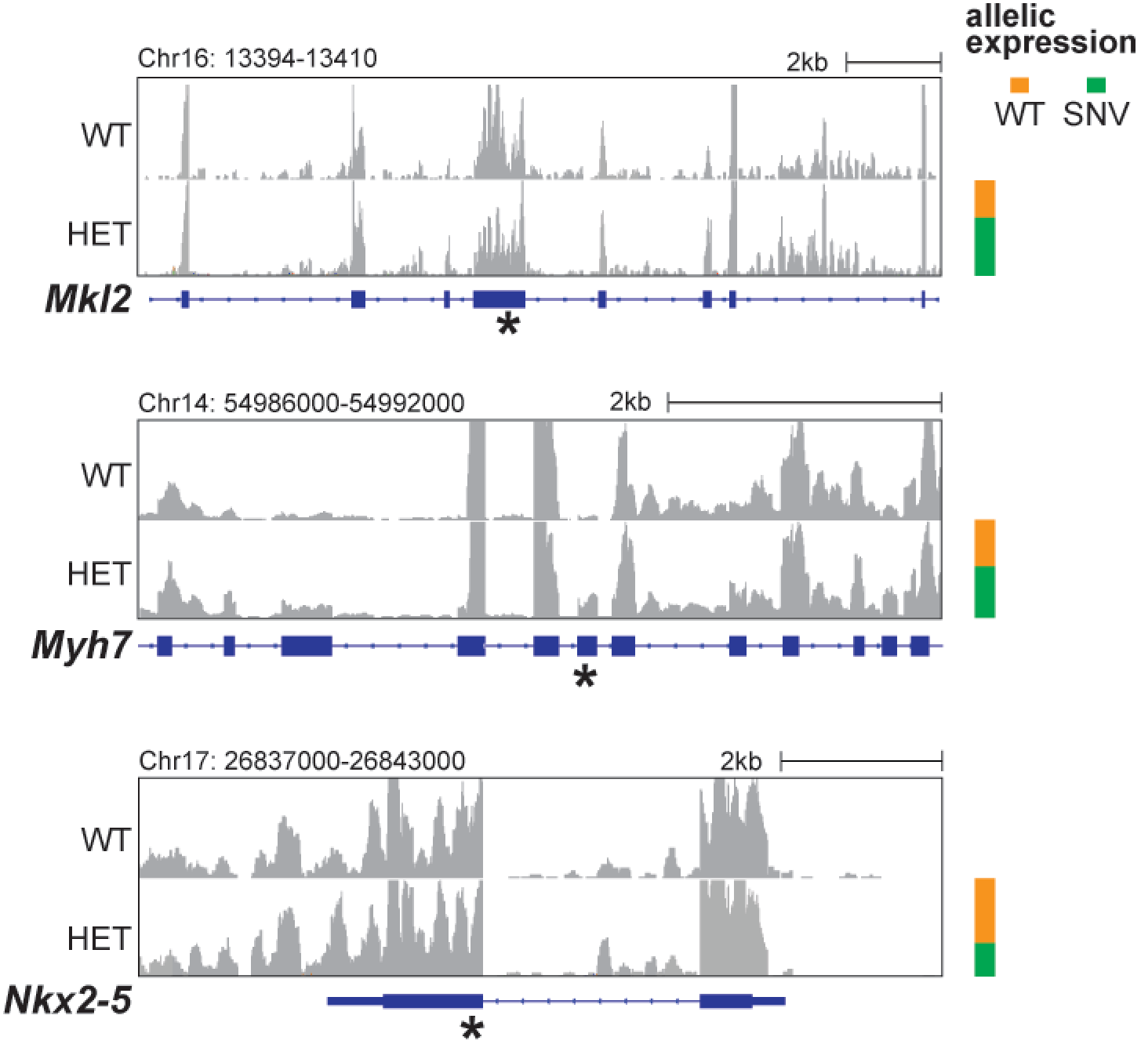
Missense variants do not disrupt mRNA splicing or allele-bias expression. Raw RNA sequencing data visualized using the Integrative Genomics Viewer from mice that encode each missense variant (*67*). The fraction of reads assigned to each allele is shown at right.

**Table S1. Exome analysis results.**

**Table S2. Primer, Cas9 guide gRNA and donor template sequences.**

**Table S3. GO term results from differentially expressed genes in Wt versus Triple Compound Heterozygous Mice.**

**Table S4. Statistically significant differentially expressed genes in Wt vs Triple Compound Heterozygous Mice.**

**Table S5. GO term results from differentially expressed genes in unaffected differentiated cells versus symptomatic LVNC.**

**Table S6. Statistically significant differentially expressed genes in unaffected differentiated cells versus symptomatic LVNC.**

